# Three-color single-molecule localization microscopy in chromatin

**DOI:** 10.1101/2024.09.09.612161

**Authors:** Nicolas Acosta, Ruyi Gong, Yuanzhe Su, Jane Frederick, Karla Medina, Wing Shun Li, Kiana Mohammadian, Luay Almassalha, Geng Wang, Vadim Backman

**Author notes:** These authors contributed equally to this work (Nicolas Acosta, Ruyi Gong).

## Abstract

Super-resolution microscopy has revolutionized our ability to visualize structures below the diffraction limit of conventional optical microscopy and is particularly useful for investigating complex biological targets like chromatin. Chromatin exhibits a hierarchical organization with structural compartments and domains at different length scales, from nanometers to micrometers. Single molecule localization microscopy (SMLM) methods, such as STORM, are essential for studying chromatin at the supra-nucleosome level due to their ability to target epigenetic marks that determine chromatin organization. Multi-label imaging of chromatin is necessary to unpack its structural complexity. However, these efforts are challenged by the high-density nuclear environment, which can affect antibody binding affinities, diffusivity and non-specific interactions. Optimizing buffer conditions, fluorophore stability, and antibody specificity is crucial for achieving effective antibody conjugates. Here, we demonstrate a sequential immunolabeling protocol that reliably enables three-label studies within the dense nuclear environment. This protocol couples multiplexed localization datasets with a robust analysis algorithm, which utilizes localizations from one target as seed points for distance, density and multi-label joint affinity measurements to explore complex organization of all three targets. Applying this multiplexed algorithm to analyze distance and joint density reveals that heterochromatin and euchromatin are not-distinct territories, but that localization of transcription and euchromatin couple with the periphery of heterochromatic clusters. This work is a crucial step in molecular imaging of the dense nuclear environment as multi-label capacity enables for investigation of complex multi-component systems like chromatin with enhanced accuracy.

## INTRODUCTION

The development of single-molecule localization microscopy (SMLM) has allowed unprecedented investigation of the nanoscopic structure of biological tissues^1–5^. These capabilities had been greatly enhanced by the capacity to image multiple molecular species, allowing the investigation of the relationship between sub-diffraction structures in space and time^6–11^. Among the most complex structures within cells is chromatin, the assembly of the genome folded into three-dimensional space^12–14^. With the rapid advancement of sequencing-based methods (chromatin immunoprecipitation sequencing, high-throughput chromatin conformation capture, etc.)^15–19^, several models describing chromatin folding such as the loop extrusion model^20–22^ have been proposed to recapitulate the findings from inferred models through studies on Topologically Associated Domains (TADs) and genome compartments^15,21–23^. SMLM studies, based on DNA-Point accumulation for imaging in nanoscale topography (DNA-PAINT) and multiplexed-fluorescence in situ hybridization^15,16^, utilized the inferred the localization of molecular labels obtained by these methods to provide information about the underlying structure of the genome upon imaging^23,24^. However, these methods rely on formamide denaturation which can result in disruption in the underlying space-filling conformations of the genome in space^33^. Therefore, it was not until the advent of chromatin electron microscopy (ChromEM)^23^, that the ground-truth for the *in vitro* sub-diffraction supra-nucleosome genome structure was observed, remained elusive at the regulatory length-scales that were beyond conventional imaging modalities (<200nm)^29,51^.

As described by Ou et al.^25^, chromatin is organized as a disordered polymer at the smallest length scales, ranging from 5 to 20 nm. Utilizing the ChromEM framework and high-angle annular dark-field imaging technique, chromatin scanning transmission electron microscopy (ChromSTEM)^26^ allows for identification of higher-order chromatin structures (packing-domains) at supra-nucleosome length-scales with a distribution of sizes and length-scales present without a characteristic partitioning into discrete heterochromatin (compacted) and euchromatin (accessible) states^27,28^. Given this surprising finding that interphase chromatin is not assembled into discrete functional territories, it is imperative to develop new methods of analysis and imaging of chromatin conformation to understand the molecular structure of supra-nucleosomal folding. At length-scales between 100-200nm, Miron et al.^29^ and Li et al.^30^ described how heterochromatin and euchromatin coupled to produce the complex formation of packing-domain (PD) structures. Using structured illumination microscopy (SIM), Miron et al. found that heterochromatin is organized within PD centers, with euchromatin and RNA-polymerase extending outward to the periphery at a label localization precision of 80-100 nm. Li et al. using a combination of single color SMLM of active RNA polymerase and partial-wave spectroscopic (PWS) microscopy^31,32^, which, although diffraction limited, is sensitive to folding of structures between 20-200 nm, found that active RNA polymerase localizes to the boundary of PDs^30^. Despite this finding, there remains a crucial gap in our understanding of the molecular assembly of the genome at length-scales between 20-100nm. Indeed, recent work utilizing ChromSTEM tomography has shown that packing domains are heterogeneous structures with variations in assembly and density at length-scales ranging from 50-200nm^33^.

Consequently, the development of multiplexed SMLM to study the spatial relationship between heterochromatin, euchromatin, and RNA polymerase at higher resolution, multi-color imaging could uncover new molecular processes in the regulation of genome structure. Multiplexed SMLM, including multi-color direct STORM (dSTORM)^9^ and (DNA-PAINT)^34^, has elucidated multiple targets and their spatial relationships in single cells with resolution up to 10 nm^34^. dSTORM utilizes immunostaining, while DNA-PAINT employs fluorophore-bound imager DNA strands that bind to its complementary docking strands attached to specific targets. Recently reported DNA-PAINT achieved labeling of up to 30 targets within cells^23^, but the denaturation-based probe binding disrupts chromatin organization by breaking down the double-strand structure of DNA with formamide^33,54^. To elucidate the chromatin domain structure while preserving its integrity, immunostaining-based super-resolution, such as dSTORM, imaging remains the only viable solution so far. However, due to the dense packing environment in chromatin, three-color dSTORM for nuclear targets has never been reported although two-color chromatin dSTORM and three-color dSTORM in cytoplasm have been documented.

Adding to the complexity of genome structure is the fact that it is inherently a disordered polymer structure, therefore its reconstruction from sparse emissions contrasts with the capacity to resolve simpler structures such as microtubules as the distribution of emission events requires more complex spatial analysis. Likewise, probe targeting of highly dense chromatin polymer with volume concentrations between 0.2 and 0.8 is challenging, which require a high label density, but the accessibility of chromatin is poor, especially to the large labels required for molecular recognition. This is compounded by difficulties in delivering high label concentrations and ensuring their diffusion through the dense nuclear environment^36–41^. Moreover, label density together with photon-budgeted localization defines spatial resolution, which is limited to twice the spatial distance between labels. Currently, the reported maximum number of colors used in chromatin super-resolution imaging remains limited to two^42–45^. Besides the prior issue, more challenges exist to imaging and analysis of chromatin at length-scales below 200 nm utilizing SMLM methods: (1) Staining protocol,imaging buffer, fluorophore stability, and antibody specificity must be optimized to achieve proper antibody conjugates. (2) Complex analysis of multiple labels is required as the relationship between euchromatin and heterochromatin with enzymes such as RNA-polymerase may be more complex than binary exclusions.

In this work, we developed a 3-color chromatin SMLM *in vitro* overcoming the challenges of multi-labeling in a dense structure by optimizing antibody conjugation timing and utilizing newly described imaging buffers^44^. To facilitate the complex analysis of 3-color SMLM, we develop a clustering-based 2-color distance analysis and 3-color joint density analysis methods to compare multiple coupled molecules within these confined spaces. We demonstrate that in contrast to prior work with 2-color chromatin SMLM showing the separation of heterochromatin and euchromatin into two partitions, three-color chromatin imaging demonstrates that the genome organizes into packing domains with euchromatin and active transcription occurring around the periphery of constitutive heterochromatin cores. This work presented a method to reliably achieve multi-color SMLM in chromatin to probe additional complex interactions with a data analysis methodology to probe the complex interactions of the genome.

## RESULTS

### Sequential labeling and optimized imaging buffer facilitate three-color single-molecule localization microscopy of heterochromatin, euchromatin and active RNA polymerases

There are multiple different approaches for targeted binding of immunofluorescent markers in super-resolution microscopy. Among the most widely used is concurrent incubation of primary antibodies against multiple targets with the subsequent addition of concurrent secondary antibodies. As each antibody is derived from a different host organism, it is generally assumed that their binding efficiencies will result in binding of the distinct targets concurrently. Therefore, we first applied the published staining protocol for 3-color SMLM labeling which was originally for Tom20, ATPB and tubulin^9^, to staining of chromatin markers associated with the differential states of genome function: H3K9me3 (constitutive, high-density heterochromatin), H3K27ac (enhancer associated euchromatin) and transcriptionally active RNA polymerase II (RNAP II)^9^. In this protocol, all three primary antibodies are incubated with the cells simultaneously during the primary antibody incubation, followed by simultaneous incubation of all three secondary antibodies during the secondary antibody incubation. Images show that all three channels are under-labeled with extremely low number of localizations (Fig. 1A Top). Although simultaneous incubation can successfully stain structures in the lower-density cytoplasm^9^, the highly dense nuclear region can non-monotonically influence molecular interactions and lead to non-specific binding, such as histone markers and active RNA polymerases. Therefore, we developed a sequential labeling method (Fig. 1B, Supplementary note), in which the cells are only incubated with one antibody at a time to achieve a higher labeling density for each target (Fig. 1A bottom, Fig. S2C). Between each labeling sequence (washing step is not noted in Fig. 1B), samples underwent repeat blocking to minimize the frequency of non-specific binding and adsorption^46^. The choice of blocking with goat serum was derived from the secondary antibody source, which blocks the attachment of Fc receptors in the sample to both the primary and secondary antibodies used in experiments^47^.

**Fig. 1.**
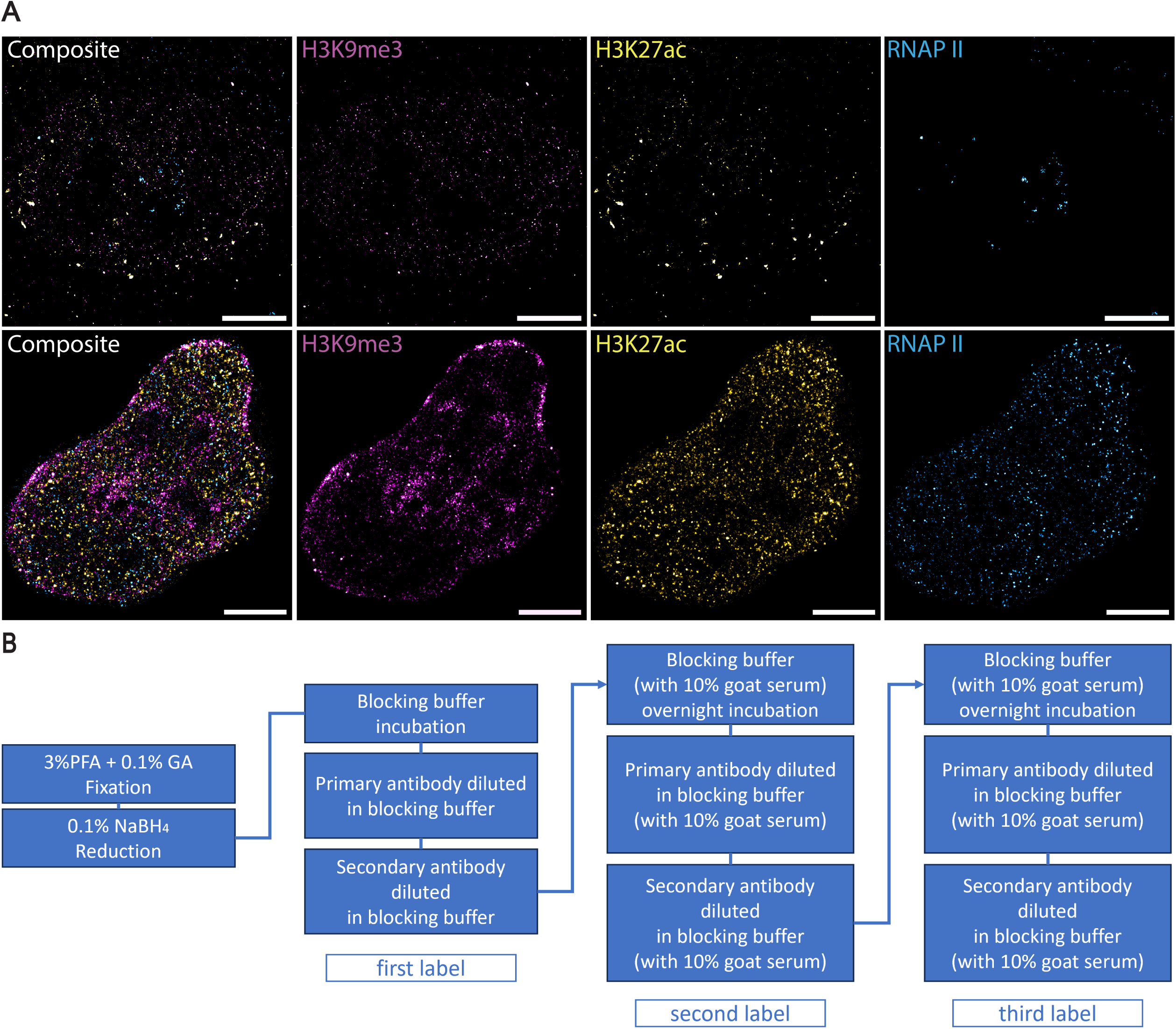
Unsuccessful and successful staining for 3-color SMLM and the flowchart for the sequential labeling. (A) Top row shows the failing staining of H3K9me3, H3K27ac, RNA Polymerase II in OVCAR5 cells, based on simultaneous staining protocol, and their merge image respectfully, while (A) bottom row displays the successful staining by sequential staining protocol in HeLa cells. All images scale bar is 5 µm. (B) describes the sequential staining protocol for 3-color multiplexed single molecule localization microscopy sample preparation.

The fluorophores we used for the 3-color SMLM were AF647, AF568 and AF488. During the imaging process, we excited the fluorophores respectively by 637 nm, 532nm, 488 nm laser lines in sequence to achieve minimum inter-channel photobleaching caused by blue light. The sequential imaging process usually took 8-10 minutes to obtain 10,000 frames for each channel. In total, one 3-color SMLM image took around 25 minutes. As a result, one limitation of multiplexed SMLM imaging is the collection of multiple cells in different conditions might take up to 4 hours or even longer. Achieving sustained emission efficiency across such a timeframe and multiple fluorophores requires careful selection of SMLM buffers. To achieve this end, we utilized a recently described optimized imaging buffer that does not rely on the catalase reduction system^46^. Despite the extended sample preparation time of 3 days for staining, this sequential imaging protocol successfully achieved 3-color SMLM for targets in the nuclear region of multiple distinct functions and molecular states.

### Simulation of different statistical distributions validate the point cloud-based and image-based methods for analysis multi-color SMLM data

With successful implementation of the 3-label immunofluorescence labeling protocol, we next developed two orthogonal analysis pipelines to investigate the higher-order structure of chromatin in multiplexed samples (Fig. 2A, Fig. S5). We pursued this strategy of two distinct analytic approaches due to the limited knowledge of the ground-truth structure of the genome. Consequently, the reliance on one modality could result in a skewed understanding of the genome based on the underlying assumptions of the analytic method. SMLM datasets can be analyzed using point-cloud based methods that directly utilize the localizations or image-based approaches that work with reconstructed images^48,49^. At present, there is no consensus or standard practice on which method provides better results; however, most analytic approaches typically employ point-cloud-based methods due to the variety of datapoint clustering analyses that exist^48,49^. Thus, we developed two analyses that report the spatial distribution of selected targets relative to a seed point to demonstrate potential differences between their results arising from their respective methodology. Each pipeline utilizes the three different targets that have been imaged in its analysis, either in a point cloud or image format. By selecting one of the labels as a seed point of interest, the relative distance of the other two targets is subsequently measured (Fig. 2A, S5). We selected H3K9me3 as our seed point due to prior studies that have investigated the organization of histone modifications via super resolution imaging showing that chromatin possess a concentric distribution of heterochromatin to euchromatin within the nucleus at the 200 nm length scale^29,30^.

**Figure 2.**
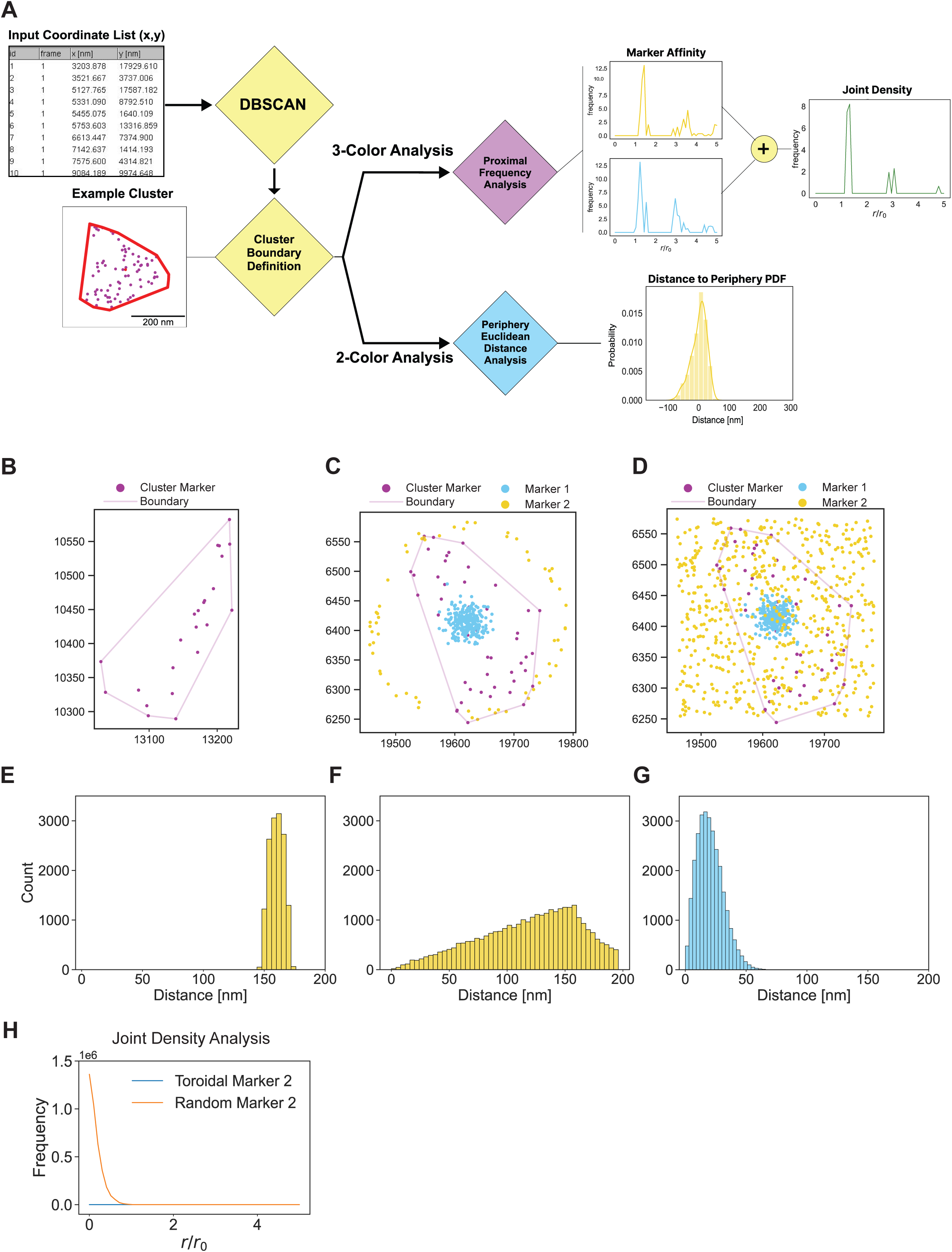
Simulated data 2-color and 3-color results for Image and Point-cloud based algorithms demonstrate algorithm robustness. **(A)** Shows our proposed pipeline for point-cloud based analysis of 2 and 3 color SMLM datasets. **(B)** Showcases an example heterochromatic cluster with fitted cluster periphery. **(C)** Shows example one of two simulated spatial distribution cases (Gaussian for Marker 1 in blue and Toroidal for Marker 2 in yellow) overlayed an example heterochromatin cluster with periphery fitted**. (D)** Demonstrates the second simulation case similar to (C) but with a random spatial distribution for simulated Marker 2 in yellow. **(E-G)** The 2-label spatial distance count histogram results for the point-cloud based algorithm with the same distributions demonstrated in (C) and (D). **(H)** Shows the joint density curves for the two test cases shown in (C) and (D) respectively, with simulation case (C) corresponding to the blue curve and simulation case (D) corresponding to the orange curve. Joint Density curve shown is relative to the heterochromatin centroid

To elucidate structure and spatial connectivity of the localized heterochromatin, we employed Density-Based Spatial Clustering of Applications (DBSCAN)^50^ which has been shown to robustly deal with different distribution. Clustering parameters epsilon and number of points considered were determined via a customized grid search algorithm which used a domain shape score to evaluate clusters and monitor clustering efficacy (Fig. S3F and Supplementary Information). For a detailed description of the grid search algorithm and our scoring method, refer to the Supplementary Information. This first step was applied to both pipelines to cluster H3K9me3 and set the seed points for either analysis (Fig. S3A). Once clustered, the dataset was used to identify their centroids via calculating the mean of coordinates of members of a given cluster and their periphery to establish landmark positions from which we can measure the spatial distance of the other two localized targets. The difference amongst these two analysis pipelines after this cluster identification is illustrated in Fig.S3, but their main distinguishing features lie in how the periphery of these H3K9me3 clusters is defined and the modality by which the spatial analysis is accomplished (image based or point cloud). In the first analysis method, the periphery is simply defined as half the max dispersion of data points assigned to an identified cluster. Thus, the periphery was a circle with a radius equivalent to half the max distance between any two points within the cluster (Fig.S5B). The second analysis invokes the Convex Hull estimation of the cluster periphery. This approach was chosen to define the boundary of our heterochromatic clusters based on localizations we observe and to avoid imposing a geometry that assumes presence of a boundary where there are no localizations (Fig. 2B).

Two approaches were proposed to investigate spatial distributions via two different inputs: image of labeled targets or coordinate point dataset of localized emission events. The image-based approach method, which from here on will be referred to as the “hybrid approach” (Fig S5.A) assumes a hybrid combination of input data which utilizes localized emission events in each channel and works with the reconstruction of these images according to the established average shifted histogram within the ThunderSTORM plugin^51^. Beginning with the DBSCAN identified clusters and their centroids, we place them in an image and then measure the distance in a pixel-wise manner from the centroids to all other pixels in the full-size image resulting in a distance cube (Fig. S5A). Using the reconstructed images of the other targets, in this case RNAP II and H3K27ac, as binary masks, we then crop and mask the distance cube slice for a given centroid to capture the positions of the other two targets within our region of interest and subsequently compile them across all clusters to establish their distribution. The second analysis method, which is referred to as the “point cloud” approach, solely uses the localization datasets. Primary clustering steps are the same as the former method, however it deviates after cluster definition (SI: Point Cloud Analysis Algorithm). This analysis calculates the Euclidean distance of targets relative to the identified cluster periphery (Fig S4A). as well as the joint density of the labeled targets (Fig. 2A). Joint density aims to uncover areas of enrichment for both other labels relative to the landmark seed points and was easily implemented in our point-cloud approach due to the format of the data (Fig. S4B).

To understand how each of these analytical methods would depend on the underlying geometry prior to applying them to real biological datasets, we tested them on simulated localization datasets that followed pre-determined statistical distributions (Gaussian, Random, Uniform, Toroidal) (Fig. SA). We employed these distributions as they are expected to be easily distinguishable from each other when measuring spatial distance to our relative seed points. In this simulated study, the seed points (cluster centroids) were determined from actual heterochromatic datasets collected to mimic observed distributions and not assume spacing between clusters *a priori* (Fig. 2B). When applying these distributions to our seed points, both the hybrid and point cloud-based methods produced visually distinct distance histograms which counted the simulated localizations into bins relative to the center of each cluster (Fig 2. E-G, Fig. S6C). While the distributions were not the same for each method, they were visually distinct depending on the distribution tested, showing that both methods can distinguish marker distributions. Similarly, we applied the joint density analysis to this simulated multi-label data where one marker was held at a constant normal distribution and the second marker was either toroidally or randomly distributed (Fig. 2C, D). In the Gaussian-toroidal case (Fig. 2C) where there is no overlap between the distributions, there is an expected flat curve within the analysis region (Fig. 2H). However, for the normal-random case (Fig. 2D) where there is direct overlap of the two markers, there is a decaying distribution with joint density expectedly decreasing farther from the landmark reference point of cluster centroids (Fig 2H). With no overlap, the curve behaves as expected with a flat distribution, however for the normal-random case there is a decaying distribution with joint density decreasing as expected the farther from the cluster centroids. The rest of this work uses the use of the point-cloud based approach as it provides more flexibility in analysis and allows for more granular spatial analysis that avoids binning when working with biological datasets.

### Enhancer associated euchromatin and active RNA Polymerases II Localize at the Periphery of Heterochromatin Clusters

Heterochromatin and euchromatin have different functional associations with the transcription of mRNA. Specifically, the presence of heterochromatin markers such as H3K9me3 within the transcription start sites (TSS) of genes is associated with their repression. Likewise, enhancer associated euchromatin, which is frequently defined by H3K27ac nucleosome modifications, are modifications to distal elements (segments tens to hundreds of kilobases away from the TSS) that when brought into contact with a gene promote its transcriptional activity. While it is hypothesized that the functional coupling of heterochromatin, enhancer associated euchromatin and active RNA polymerase can be inferred from their spatial relationships, prior to this work it has not been demonstrated with required resolution to uncover their interactions. Therefore, we utilized our multiplexed SMLM imaging and analysis approach to study the spatial relationship between heterochromatin (H3K9me3) and enhancer-associated euchromatin (H3K27ac), and between heterochromatin (H3K9me3) and RNAPII. After the validation of the algorithms on simulated dataset, we apply the 2-color distance analysis on H3K9me3, H3K27ac and RNAPII SMLM data from HeLa cells. Heterochromatin localizations are clustered by DBSCAN with parameters *r* = 50 nm and *pts* = 3. The heterochromatin cluster’s size is calculated as the polygon size of the cluster’s convex hull. The reported average size of DNA packing domain is 80 nm in radius^26^. Owing to the distribution in sizes of packing domains, we partitioned heterochromatic clusters based on their effective radii which is defined as the radius of the circle that inscribes the convex hull fit of the cluster datapoints. Clusters were split into small (25-40 nm), medium (40-80 nm) and large domains (80-250 nm) based on their sizes, defined as the radii of the circle with the same size as the convex hull of the clusters. To investigate the locations of surrounding euchromatin and RNAPII, a search radius of 1.5 times of the heterochromatic cluster size (radius) is used for distance calculations. The coordinates of each domain’s center are determined by averaging those of convex hull vertices of the cluster (Fig. S4A). To quantify the distance of each euchromatin or RNAPII localization within the searching radius of a heterochromatic domain to the domain periphery, we draw a line connecting the localization point to the domain center (Fig. S4A). The distance from the point where this line intersects the convex hull to the localization point is defined as the distance to the periphery (Fig. S3). If the localization point is outside the heterochromatin domain, the distance to periphery is positive, otherwise it is negative. We found that H3K27ac and RNAPII are generally localized near the heterochromatic domain periphery. In large domains, H3K27ac is closer to the heterochromatic domain center compared to RNAPII, whereas the opposite is observed for medium and small domains (Fig. 3). For large heterochromatin domains: H3K27ac meansdistance average distance to periphery is -1 nm, RNAPII meansdistance average distance is 8 nm (Fig. 3C, F). For medium heterochromatin domains: H3K27ac average distance is 1 nm, RNAPII distance average distance is -2 nm (Fig.3B, E). For small heterochromatin domains: H3K27ac average distance is 3 nm, RNAPII average distance is -2 nm (Fig.3A, D). In conclusion, within the 1.5x searching radius of a H3K9me3 cluster, the average distance of H3K27ac and RNAPII localizations to the periphery is close to zero, indicating a closed structure of heterochromatic domain surrounded by euchromatin and RNAPII. However, the paired distance analysis only elucidates the spatial relationship between euchromatin and heterochromatin or RNAPII and heterochromatin, not providing information on the interaction among all three markers, e.g. where the coupling of euchromatin and RNAPII is regarding the heterochromatic domain.

**Fig. 3.**
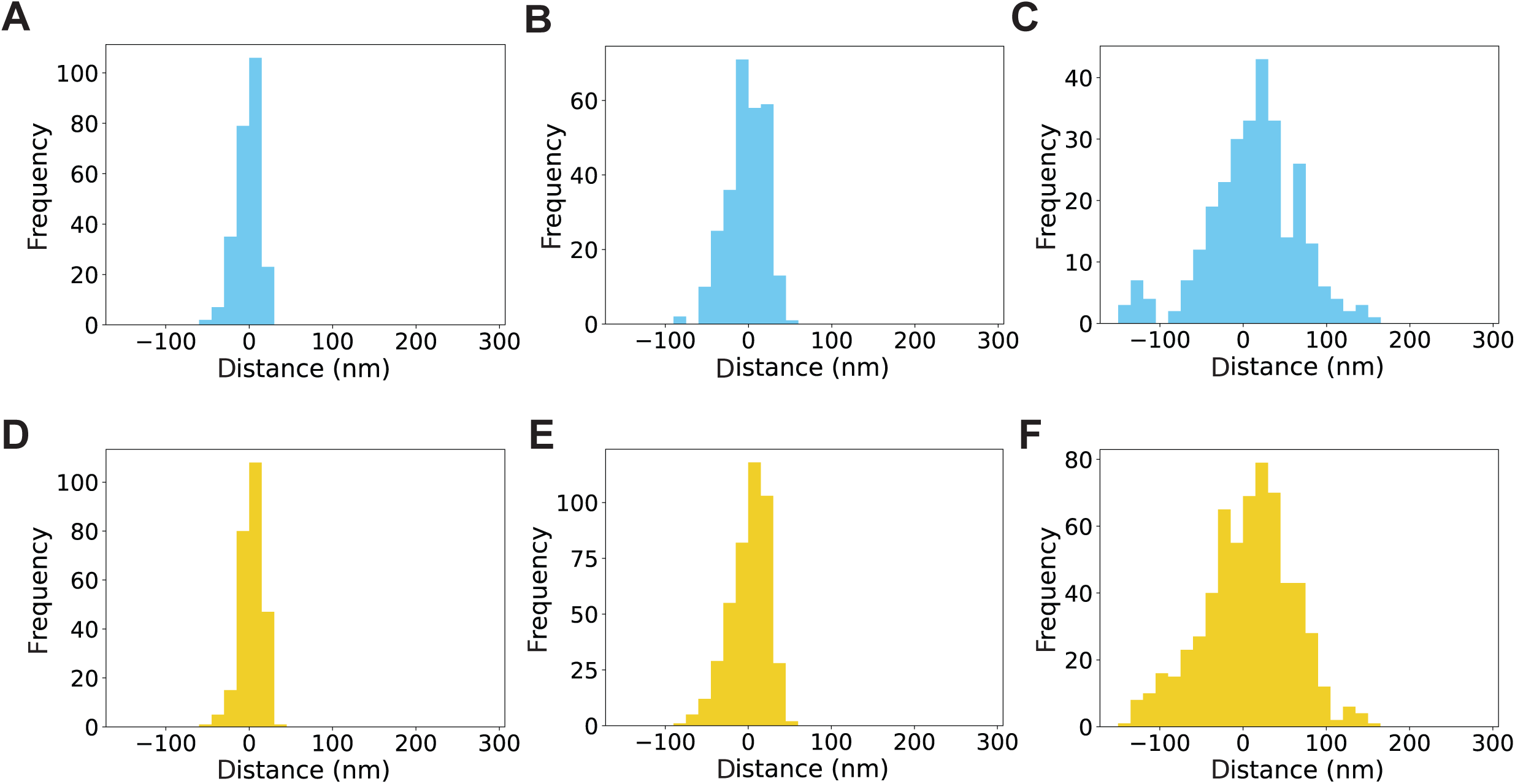
2-color distance analysis between heterochromatin and euchromatin, heterochromatin and functional RNA polymerase for different size of heterochromatic clusters. **(A-C)** 2-color analysis: distance of RNAPII localizations relative to heterochromatin cluster periphery for small (A), medium (B), and large clusters (C) respectively. **(D-F)** Showcases the 2-color analysis results for H3K27ac localizations relative to heterochromatin cluster periphery for small (D), medium (E), and large clusters (F) respectively

### Three-Color SMLM joint analysis reveals spatial coupling of heterochromatin, euchromatin and RNA Polymerases

The spatial organization of chromatin and transcription within the nucleus presents a complex landscape of functional interactions. While traditional models posit distinct compartmentalization of heterochromatin, euchromatin, and transcription sites, emerging evidence suggests a more intricate arrangement^26,29,30^. We hypothesized that transcription occurs preferentially at the interface of heterochromatic domains, rather than in spatially segregated regions. To investigate these relationships and compare the organization of heterochromatic and euchromatic regions, we developed a three-color joint analysis method for multi-label single molecule localization microscopy (SMLM) data. This approach enables the examination of spatial relationships between chromatin components and transcription sites, providing insights beyond those obtainable from conventional two-color analyses. Our approach begins by identifying heterochromatin clusters as seed points. For each cluster, we define a search area extending five times the cluster’s length while maintaining its morphology as defined by convex hull fitting (Fig. S4D). Within this search zone, we locate nearby euchromatin (H3K27ac) and RNA Polymerase II (RNAP II) localizations. We then calculate their distances from the domain center and normalize these distances by the search radius, yielding normalized radial density distributions for both euchromatin and RNAP II. With these points identified, we examine the affinity of each individual centered target (RNAP II or H3K27ac) for the other. We calculate normalized radial density distributions for two cases: centering RNAP II and searching for H3K27ac (Fig. 4E), and the inverse case of centering H3K27ac and searching for RNAP II (Fig. 4F) within heterochromatic clusters. These distributions are then used to compute the joint density shown in Fig. 4G by taking the geometric mean of both cases (Supplementary Information). To account for localization uncertainty in the SMLM data, we applied this 3-color joint density analysis only to large heterochromatic domains defined in the previous section. We validated our algorithm using simulated data (Fig. 2H), ensuring its reliability for subsequent analyses of spatial coupling between chromatin states and transcription sites.

**Fig. 4.**
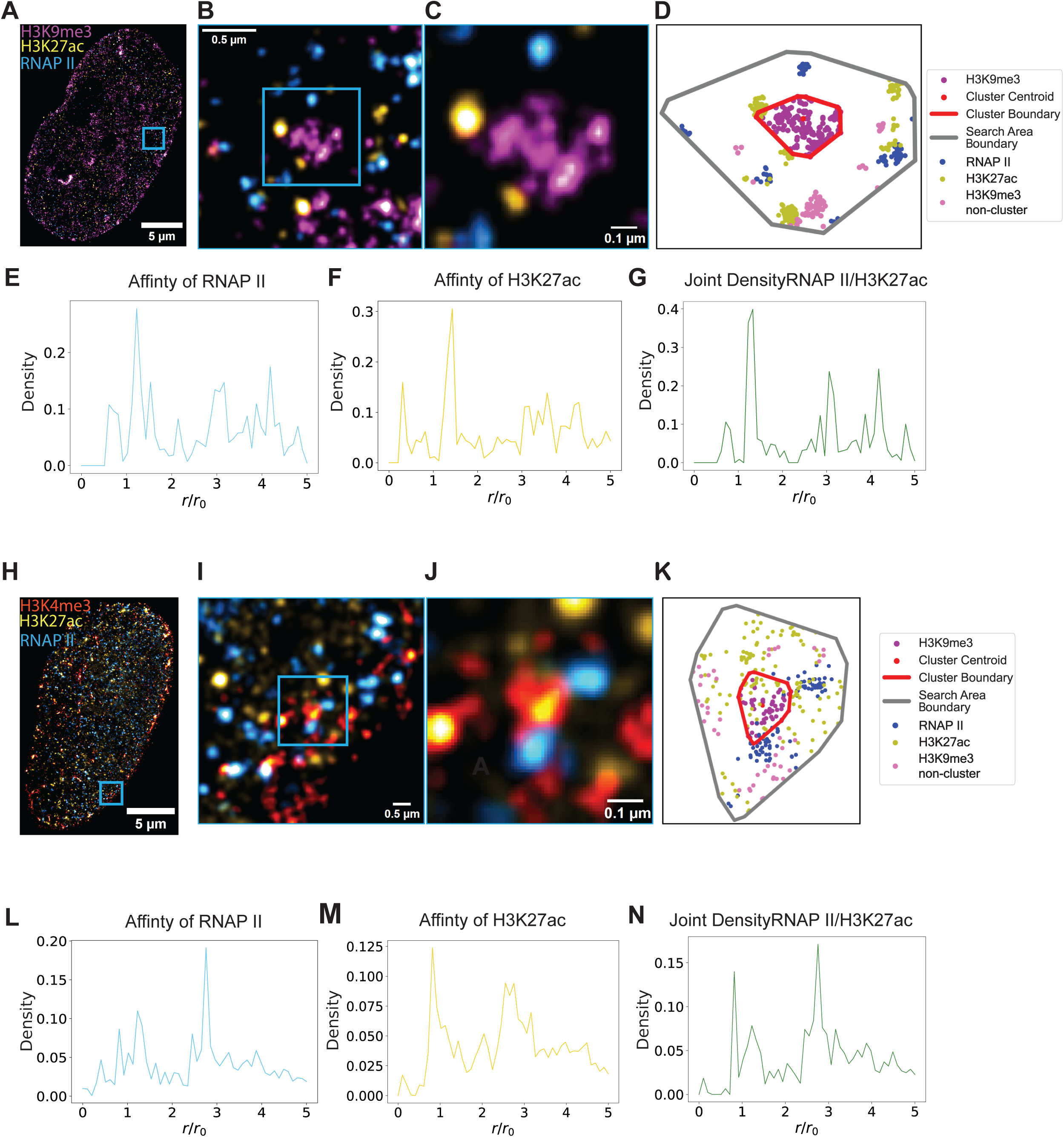
Joint Density Analysis of Multi-label SMLM Imaging in HeLa cells reveals co-existence of RNAPII and H3K27ac on Heterochromatic cluster periphery but overlap in Euchromatic clusters. **(A)** shows the three-color SMLM image of a HeLa cell labeled for H3K9me3 (magenta), H3K27ac (yellow) and RNA polymerase II (blue). **(B)** and **(C)** are the highlighted blue insets in the images demonstrating the highlighted heterochromatic cluster with peripheral RNAPII/H3K27ac. **(D)** Is the an example cluster from localization dataset for inset shown in (C) plotted. Red Boundary represents periphery of heterochromatic cluster; Gray boundary represents boundary of analysis region of interest, **(E/F)** are the joint density of RNA polymerase II to H3K27ac and affinity of H3K27ac to RNA polymerase II regarding H3K9me3 clusters respectively across n=4 HeLa cells. **(G)** shows the joint density for heterochromatic clusters, combining data from (E) and (F). **(H)** shows three-color SMLM image of a Hela Cell labeled for H3K4me3 (red), H3K27ac (yellow) and RNA polymerase II (blue). **(I/J)** are the highlighted blue inset demonstrated in **(H)** with **(K)** being the example localizations shown in G plotted with the red boundary being the periphery of H3K4me3 cluster and gray the analysis region of interest boundary. **(L/M)** are the joint density of RNA polymerase II to H3K27ac and affinity of H3K27ac to RNA polymerase II regarding H3K4me3 clusters respectively across n=4 HeLa cells. **(N)** shows the joint density for euchromatic clusters, combining data from (L) and (M).

Using this method, we examined the spatial organization of enhancer-associated chromatin (H3K27ac), active transcription sites (RNAP II), and heterochromatin domains (H3K9me3). Fig. 4 A-D shows an example HeLa cell labeled with these three markers, visually showing how euchromatin and active RNA polymerases localize around a representative heterochromatic cluster. The normalized radial density distribution of RNAP II localizations proximate to H3K27ac is shown in Fig. 4E, while Fig. 4F presents the inverse case. Fig. 4G displays the joint density distribution. Our analysis revealed that the largest peak of the joint frequency distribution for H3K9me3 domains is located outside the normalized cluster size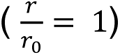, indicating that the most frequent coupling of H3K27ac and RNAP II occurs outside the heterochromatic domains. This finding supports the hypothesis that transcription takes place at the interface of heterochromatic clusters. To compare this organization with euchromatic clusters, we performed a similar analysis using H3K4me3 as a marker for promoter-associated euchromatin. Fig. 4H-K shows a HeLa cell labeled with H3K4me3, H3K27ac, and RNAP II. The distribution results for H3K4me3 clusters are presented in Fig. 4L-N. In contrast to the H3K9me3 results, we observed that the coupling of H3K27ac and RNAP II occurs inside the euchromatic (H3K4me3) clusters. These findings challenge the concept of transcription factories and suggest a new model of nuclear organization: heterochromatin domains serve as organizational hubs, with euchromatic regions and active transcription sites preferentially localized at their interfaces. This three-color joint density analysis uncovers a hierarchical chromatin architecture that orchestrates transcriptional activity, providing insights into nuclear function that extend beyond the capabilities of conventional two-color SMLM approaches.

## DISCUSSION

This study introduces the first reported method for three-label single-molecule localization microscopy (SMLM) targeting chromatin in the dense nucleus, achieving uncertainty of ∼15-20 nm. Our method enables simultaneous visualization and quantification of multiple chromatin markers, providing insights into the higher-order structure of the genome at the supra-nucleosomal length scale (10-200 nm). This technique, combined with our developed analysis methods, reveals a more nuanced understanding of chromatin organization, challenging previous models and opening new avenues for investigating nuclear architecture and functions. With this approach we observed cohesive, domain-centered structure rather than spatially distinct groups of chromatin states, and the preferential localization of euchromatin and active RNA polymerase II around heterochromatic clusters. The cornerstone of our approach is an optimized sequential labeling protocol, enabling robust labeling of three targets within the dense nuclear environment yielding higher localization density compared to previously published methods^9^ (Fig. 1B, SI Figure S2C). This innovation, combined with our three-label joint analysis (Fig. S4 D-F), allows for the co-registration of three distinct chromatin states or protein localizations, offering a more comprehensive view of nuclear organization than previously possible with two-color or diffraction-limited imaging techniques.

To ensure widespread applicability and reproducibility of our technique, we developed a protocol that is both powerful and accessible. This protocol employs commercially available antibodies and modified blocking buffers (SI Table 1), making it accessible and easily implementable across various cell lines. We employ modified blocking buffers that include the serum of the host species of our secondary antibodies, used after every target labeling stage. This secondary and tertiary blocking step helps improve target specificity and ensures that all residues for potential off-target binding are blocked before adding our specific primary antibody for the desired targets. Furthermore, overnight incubations at 4°C are performed to ensure proper labeling. Our protocol is sufficient and optimized for various cell lines used regularly (Fig. 1A, Fig. S2), further promoting the ease of implementation for multiplexed labeling of targets within nuclei. While we have established a robust process for targeting 3 structures in the nucleus, there is a limit to the number of available targets, as each target must use antibodies that have no crosslinking capacity. This obstacle is potentially overcome by fluorophore-conjugated primary antibodies, but this will sacrifice blinking density, thus reducing resolution. To achieve multi-color SMLM with more than 3 immunofluorescence labels, more laser lines should be used, or spectroscopic SMLM should be introduced to distinguish different fluorophores excited by the same laser line. The development of this technique required overcoming several challenges inherent to imaging the dense nuclear environment. Unlike the cytoplasm or cellular membrane, where multi-label SMLM has been previously applied, the nucleus presents a unique set of difficulties due to the proximity of targets and their relatively smaller size (<20 nm) compared to the two-antibody systems used to label them. Our sequential labeling approach, combined with optimization of blocking and incubation steps, allows us to achieve specific labeling of multiple targets in this challenging environment. Moreover, our technique addresses limitations of previous chromatin imaging methods. While recent work using structured illumination microscopy (SIM) and Lattice Light Sheet Microscopy has provided valuable insights into chromatin organization^10,23,29,42^, these methods are limited by their lateral resolution (>100 nm) and label number. Our approach, with its higher lateral precision of labeled targets, allows for a more detailed examination of chromatin organization at smaller scales, capturing more structural details of this complex organization.

Fully leveraging our localization precision and three-label data, we developed novel analysis methods tailored to the complexities of chromatin structure at the packing domain length scale (<100 nm)^26^. Our point-cloud based method, combined with a Convex Hull fitting algorithm, allows us to base our analysis on actual localizations within each cluster, avoiding bias from assumed shapes. This approach is particularly important given the absence of ground truth morphology for chromatin structures at these length scales. By fitting a polynomial to the cluster periphery based on available points (Fig. S4A), we can more accurately define the boundaries of chromatin domains without imposing assumptions of circular or ellipsoidal shapes (Fig. S5B). Additionally, our three-color joint density analysis provides insights into the spatial joint occupancy of different chromatin markers relative to specific landmarks. This analysis method not only makes use of the three-label dataset but also allows us to determine the distribution of these modifications together. By exploring the affinity of these marks for each other relative to a reference point (Fig. S4D-F), we can quantitatively validate canonical relationships, such as transcription occurring in transcriptionally active chromatin at the supra-nucleosome level, while also elucidating the actual distribution. This level of analysis is only possible with three-label nuclear SMLM imaging capabilities combined with proper analysis to explore the data as we cannot make claims of affinity and joint occupancy from paired datasets. Furthermore, our technique allows for the investigation of size-dependent effects on chromatin organization. By partitioning clusters into different size groups, we can observe how the spatial relationships between different chromatin marks change as a function of domain size (Fig. 3). This analysis reveals complexities in chromatin organization that were not apparent in previous studies using lower-resolution techniques or two-color imaging^43^.

Our analysis of the spatial relationships between H3K9me3 (constitutively repressed chromatin), H3K27ac (enhancer-associated chromatin), and RNAPII (active transcription) reveals a cohesive, heterochromatin domain centered structure. This finding challenges the traditional view of chromatin as being organized into distinct “active” and “inactive” compartments, suggesting instead a more nuanced and interconnected organization. When applied to biological distributions of histone modifications and proteins in HeLa cells, both image-based and point-cloud based spatial analysis revealed a preference for euchromatic and RNAPII to localize at the periphery of heterochromatin clusters (Fig.3, Fig. 4E-G, Fig. SD). This observation supports the idea of a functional organization of chromatin, where transcriptionally active regions are positioned at the interfaces between different chromatin domains, potentially facilitating rapid transitions between active and inactive states. Interestingly, we found a dependence on cluster size for the resultant distributions. When clusters are partitioned into three groups (Small <40 nm, Medium 40-80 nm, Large: 80 nm-250), the distributions all center around zero, but from small to large, we observed a shift from a single peak normal-like distribution to a potentially multimodal distribution with a wider range of values (Fig.3). This size-dependent organization suggests that chromatin domains may have different functional properties depending on their size, with larger domains potentially harboring more complex internal structures. Our three-color joint frequency analysis showed that heterochromatic clusters have a peak joint density on their periphery, while euchromatic clusters have a peak joint frequency within (Fig. 4 E-G/L-N). This indicates a joint occupancy distribution rather than a layered one, providing a more nuanced understanding of chromatin organization. The observation that active transcription (RNAPII) and enhancer-associated chromatin (H3K27ac) frequently co-localize at the periphery of heterochromatic domains suggests a model where transcriptionally active regions are positioned to allow rapid access to regulatory elements while maintaining overall chromatin compaction. These findings have important implications for our understanding of gene regulation and nuclear organization. The joint occupancy distribution revealed by our three-color analysis challenges simplistic models of chromatin organization and highlights the importance of considering multiple chromatin states simultaneously. Our findings suggest that the nuclear landscape is better described as a continuum of intermingled chromatin states rather than discrete compartments, with functional elements like enhancers and active polymerases positioned at the interfaces between different chromatin domains. The ability to simultaneously visualize multiple chromatin states and protein localizations allows for a more comprehensive analysis of how nuclear organization is remodeled during these processes. Future studies using our technique could provide valuable insights into the mechanisms underlying cellular plasticity and the role of chromatin reorganization in disease development.

The integration of our imaging protocol and associated analysis opens exciting avenues for future research. This approach can be extended to study chromatin reorganization during various cellular processes, such as cell differentiation, cell cycle progression, and in response to external stimuli. By comparing chromatin structure in different cell types or disease states, we can gain insights into the role of chromatin architecture in maintaining cellular identity and in the development of pathological conditions. The ability to simultaneously visualize multiple chromatin marks and protein localizations opens new possibilities for studying the dynamics of gene regulation. Future adaptations of our technique for live-cell imaging could provide unprecedented insights into how chromatin organization changes in real-time during processes such as transcription activation or cellular differentiation. Combining multi-color SMLM imaging and advanced spatial analysis algorithms enables the extraction of meaningful information from complex biological datasets and generates testable hypotheses for future studies. Our work not only enhances our understanding of chromatin architecture but also provides robust methodology for future investigations into the interplay between chromatin structure, gene transcription, and epigenetic processes.

## MATERIALS AND METHODS

### Cell culture

HeLa cells were grown in RPMI 1640 medium (Thermo Fisher Scientific, Waltham, MA; catalog number 11875127). The medium was enriched with 10% fetal bovine serum (Thermo Fisher Scientific, Waltham, MA; catalog number 16000044) and penicillin-streptomycin (100 μg/ml; Thermo Fisher Scientific, Waltham, MA; catalog number 15140122).

### Imaging buffer preparations

DABCO [1,4-Diazabicyclo-(2.2.2)-octane] (D27802, Sigma) was dissolved in distilled water to prepare a 1M stock solution, with 12M HCl added until the powder was completely dissolved, and the pH reached 8.0. Maintaining this pH is crucial as it primarily determines the pH of the overall buffer. The stock solution can be stored in the refrigerator, in the dark, for several weeks. DTT 1M (43816, Sigma) was used directly as purchased and stored in the refrigerator for several weeks. Sodium sulfite (S0505, Sigma) was dissolved in 10× PBS to a concentration of 1M and can be kept at room temperature on the bench for several weeks. Typically, we prepared 40mL of buffer at a time, adjusting the pH with NaOH and HCl using a pH meter, and then stored it in the refrigerator.

### 3-color SMLM Sample preparation

Primary antibodies rabbit anti-H3K9me3 (Abcam), mouse anti-H3K27ac (Thermo-Fisher), rat anti-RNA Polymerase II (Abcam) and mouse anti-H3K27me3 (Abcam) were aliquoted and stored at –20°C. Secondary antibodies goat anti-rabbit AF647 (Thermo-Fisher), goat anti-rabbit AF568 (Thermo-Fisher), goat anti-rat AF488 (Thermo-Fisher) were stored at 4°C. For catalog numbers and buffer conditions see supplementary information.

The 3-color SMLM sample preparation has 3 sequential staining processes for 3 targets.

1. The cells were plated on No. 1 borosilicate bottom eight-well Lab-Tek Chambered cover glass with 12.5k cells in each well. After 48 h, the cells were fixed in 3% paraformaldehyde and 0.1% glutaraldehyde in PBS for 10 min, and then washed with PBS once, quenched with freshly prepared 0.1% sodium borohydride in PBS for 7 min, and rinsed with PBS three times at room temperature.
2. The fixed samples were permeabilized with a blocking buffer (3% bovine serum albumin (BSA), 0.5% Triton X-100 in PBS) for 20 min
3. Samples were incubated with rabbit anti-H3K9me3 (Abcam) in blocking buffer for a minimum of 2 hours at room temperature and rinsed with a washing buffer (0.2% BSA, 0.1% Triton X-100 in PBS) three times.
4. The fixed samples were further incubated with the corresponding goat secondary antibody–dye conjugates, anti-rabbit AF647 (Thermo Fisher), for 40 min, washed thoroughly with PBS three times at room temperature and stored at 4°C. Upon this step, the staining for the first target (H3K9me3) is finished.
5. Sample is then incubated overnight in a modified blocking buffer with goat serum for the following 2-color and 3-color staining (see Supplementary Information).
6. After overnight blocking step, the sample would go through the same process as Step 2-3, however with modified blocking and washing buffers (see Supplementary Information). The second target primary antibody, in this case is a rat anti-RNA Polymerase II (Abcam) and the secondary antibody is goat anti-rat AF488 (Abcam).
7. Repeat steps 5-6 for third target (overnight blocking, then primary and secondary antibody incubations) for the third target with primary antibody of mouse anti-H3K27ac (Thermo Fisher) and the secondary antibody is goat anti-rat AF568 (Abcam). Once the secondary antibody step is done, wash two times in PBS and store samples at 4 degrees centigrade if not imaging immediately.

### SMLM System Configuration and Imaging

The STORM optical instrument was built on a commercial inverted microscope base (Eclipse Ti-E with the perfect focus system, Nikon). 2 continuous lasers were used for illumination, one of which has 5 different wavelengths (405 nm, 488 nm, 532 nm, 552 nm and 637 nm, OBIS Laser Box, Coherent) while the other is a green laser with 532 nm emission (MGL-FN-532, Changchun New Industries Optoelectronics Tech. Co., Ltd.). The lasers are collimated through a with an average power at the sample of 3 to 10 kW/cm3. Images were collected via a 100× 1.49 numerical aperture (NA) objective (SR APO TIRF, Nikon) and sent to an electron-multiplying CCD (iXon Ultra 888, Andor). At least 10000 frames with a 30 ms or 50 ms acquisition time were collected from each sample for each wavelength channel. The samples labeled with H3K9me3 or H3K4me3 (AF647), H3K27ac (AF568), RNA Polymerase II (AF488) will be firstly imaged under 647 nm laser and then 532 nm laser, lastly 488 nm laser to avoid inter-channel photobleaching caused by short wavelength light.

## DATA AVAILABILITY

All data generated or analyzed during this study will be made available upon reasonable request to the corresponding author. This includes raw image files, processed data, and analysis scripts.

## ACKNOWLEDGEMENTS

This work was supported by NIH grants U54CA268084, U54CA261694, and R01CA228272, National Science Foundation grant EFMA-1830961, and CBET-2430743, and philanthropic support from Rob and Kristin Goldman, Mr. David Sachs, and the Christina Carinato Charitable Foundation.

## SUPPLEMENTARY FIGURE TITLES AND LEGENDS

**Figure S1. Schematic of three-color single-molecule localization microscopy**

All the data were collected on this Eclipse Ti-E NIKON microscope. 3 labels (AF647, AF568 and AF488) were excited by laser line 637 nm, 532nm, 488 nm respectively.

**Figure S2. Example three-color SMLM images from BJ Fibroblast and HCT116 showcase functionality of labeling protocol in various cell lines**

**(A)** Showcases individual channels (H3K9me3: magenta, H3K27ac: yellow, and RNAPII: cyan) and composite image of all three channels in BJ fibroblasts. **(B)** Showcases the same as in A but in HCT116 cells. All scale bars are 3 µm. **(C)** Localization density for each labeling protocol. We observe a higher labeling density for our sequential labeling protocol (HeLa n=4) than in the combined labeling case (OvCar5 n=1).

**Figure S3. Domain Shape Score parameter search method and Example partitions of DBSCAN identified heterochromatic clusters**

**(A-C)** Identified heterochromatin (H3K9me3) clusters from DBSCAN with epsilon = 50 and minimum number of samples = 3 for (small <40 nm), medium (40-80 nm) and large (80-250 nm) clusters respectively. **(D)** Example ellipse fitting method used in Domain Shape Score algorithm. Sample points with fitted periphery via Convex Hull method in black with fitted ellipse in dashed red. **(E)** Histogram of Domain Shape Score values for all identified clusters in a single nucleus example for epsilon =50 and minimum number of samples = 3. **(F)** Parameter grid search for using Domain Shape Score as optimizing parameter. (50,3) provided the highest score for heterochromatin (H3K9me3) dataset indicating proper parameters for subsequent cluster analysis.

**Figure S4: Two Marker analysis and Three marker analysis diagrams demonstrating counting strategy for multi-label SMLM analysis algorithms.**

**(A-C)** Pictorially demonstrate the steps for two color SMLM analysis. The first panel demonstrates periphery fitting of DBSCAN identified cluster. B demonstrates the analysis region definition which entails a 1.5 * r0. (C) demonstrates distance calculation which measures distance in the direction of mark relative to the centroid of a cluster but subtracts the radial distance of the vector from center to periphery boundary. Only remaining distance measured relative to periphery with directionality considered. **(D-F)** Pictorially demonstrates the steps for three color SMLM counting. (D) is the same as (B) however the analysis region for three color SMLM is 5 times the radius of the cluster. (E) Is the affinity of Marker 2 (yellow) to Marker 1(Cyan). This is done by counting the number of times there is a marker 2 present within 100 nm of marker 1. (F) is the same as (E) but instead of centering marker 1, marker 2 is centered and we count affinity of marker 1. This counting method is used to generate the joint density affinity plots shown in Figure 4. **(G-H)** demonstrate how the area for 3 color analysis is partitioned for Joint Density plots. (H) Demonstrates that partitioning into 50 bins such that every step of r0 (1*r0,2*r0…) has 10 bins between. (J) is a zoom showing the partition. 1 delineates the red periphery line of the cluster and 5 is the outer most boundary.

**Figure S5: Proposed Analysis pipeline for Image based approach for multi-label SMLM**

Figure demonstrates the analysis for 2-color analysis for H3K9me3 and RNAPII. The premise of this approach is to use H3K9me3 as seed points for analysis and uses the centroid of the identified DBSCAN clusters for analysis reference points. Subsequently, a distance cube, where each slice corresponds to an identified cluster is used to calculate all possible distances relative to the heterochromatic cluster. We use a binarized image of the second target. In this case the RNAP II, to mask the distance cube for a given cluster and show the remaining distance pixels. Pixels across all clusters are concatenated and then used as dataset for analysis for a given mark. The goal is to understand the spatial distribution of a given mark relative to seed reference point.

**Figures S6: Simulated distributions and their image counterparts as inputs for Image based 2-label SMLM analysis demonstrate algorithm robustness.**

**(A)** Scatter plot of simulated distributions for Gaussian, Random, Toroidal and Uniform test cases. Distributions shown are the sum of all sampled points such that the resultant distribution across all clusters follows a given spatial distribution. **(B)** Image based counterparts for simulations shown in (A). Pixel size in the image is 26 nm. Similar to (A) the image represents the sum of all samples that were deposited around cluster seed points to have trend that follows the given spatial distribution. **(C)** Count Histogram results for spatial Euclidean distance measurements of given distribution relative to heterochromatin cluster center. Gaussian, Random Toroidal and Uniform test cases resulted in visually distinct distributions. **(D)** Kernel Density estimations for distance distributions of RNAP II and H3K27ac relative to H3K9me3 clusters in HeLa Cells (n=4, clusters = 5000) using same image-based method describe in Fig S5. Distance is measured relative to the periphery and shows similar centering near the periphery of these heterochromatic clusters.

## SUPPLEMENTARY METHODS AND MATERIALS

### Algorithm Descriptions and Considerations

The motivation and explanation of each algorithm is detailed below. For a pictorial depiction of each algorithm pipeline, see Supplementary Figure 5.

#### Image Based Analysis Algorithm

Image based analysis algorithm, or the hybrid analysis algorithm, (Fig. S5A) relies on the DBSCAN method from the scikit-learn Python library to cluster the heterochromatic marks. Circles are used to describe the geometrical properties of each cluster due to its simplicity and we acknowledge that this assumption creates a limitation in identifying the actual shape of a cluster. For each cluster, the center of the circle is chosen to be the geometric centroid of the cluster points, and its effective diameter is defined by the maximum Euclidean distance between all point pairs. Yet since the cluster points fall within the pixel size of our reconstructed images (26 nm), a image based segmentation of the border of our heterochromatic clusters is not possible, thus this approach was taken. After determining the clusters, their centroids and their boundaries, we create distance cube (X, Y, N) where X, Y are the dimensions of the reconstructed image as defined as 5 times the size of the size of our ROI during acquisition, and N was the number of clusters identified after size filtering. Filtering for size was done to ensure no clusters that were beneath our precision or too large to be biologically relevant were considered in the analysis. These thresholds are the same as mentioned in the main text. Each slice in the distance cube corresponds to each cluster such that slice indexing (0,1,2, 3,…,N) correspond to clusters indexing (0,1,2,3,…,N). This was done to easily keep track of each cluster in visual format. This distance cube, as shown in Fig. S3A, has values of each pixel relative to the centroid of that given cluster. For example, the first slice is based on the cluster (index 0) has distance values corresponding to its centroid (x0, y0) and the next slice which has a different cluster centroid (index 1) will base its distance values of (x1, y1). The logic behind this was to calculate all distance values once and not repeatedly calculate distance each time for each cluster. With a distance cube that contains the respective pixel distances to each identified cluster, we then used the reconstructed images for the other two labeled targets, in this case RNAPII and H3K27ac, and used binary thresholding to keep pixels with values greater than 250. Using a set threshold was done to ensure only the brightest pixels since it is assumed that they have similar target values. This selection would allow us to count high occurrences of our labeled targets in our distribution analysis. We use the average shifted histogram algorithm that is pre-built into the ThunderSTORM Image J plugin to reconstruct our images. Since this is essentially binning the localizations into respective bins, the brighter the pixels indicate the more estimated localization datapoints in that bin, as such we want to use a set threshold for our spatial distance analysis. This ensures that we consider a similar number of events when measuring distance relative to our clusters. With binarized images, we then mask the distance cubes such that all remaining pixels in the distance cube slice would correspond to enriched areas of our markers. The remaining pixels are then recorded and then adjusted either to report distance to centroid (no change) or distance to periphery (distance – cluster size). All distances were concatenated into one array across all imaged cells included in analysis and then processed to generate distance histograms and kernel density estimation of distance to arrive at the distribution of distance of our markers (RNAPII and H3K27ac) relative to our heterochromatic clusters.

#### Point Cloud Analysis Algorithm: Paired Distance and Joint Density measurements

This analysis method uses the points identified via Thunder-STORM directly and as such is referred to as point-cloud, since the scatter of points often resembles a cloud of particles. Point-cloud algorithms used in this study started with the same DBSCAN clustering method but differ on use of clustered data. Cluster boundary was defined either by fitting using the scikit-learn Convex Hull method to fit a polynomial to the external cluster points. This approach was taken as it based our reference point, the periphery, on data that was acquired and made no assumptions on the morphology which was a limitation of the previously described image-based approach. This cluster boundary is the reference point for both the Euclidean distance measurements to the other targets (2-color paired analysis) and the Joint density analysis (3-color analysis). To start, the identified periphery was used to define an analysis region of interest by doing a 1.5*scaling of the original vertices of the convex hull defined cluster periphery. Please see the following section for the justification of this scaling value and its use in our analysis. Distance to our labeled targets, RNAP II and H3K27ac, was measured relative to the periphery by drawing a vector from the centroid of the cluster to the localization (RNAPII/H3K27ac) in question. The distance from the centroid to the cluster boundary is then measured along that same vector and the difference of these measurements is assigned to that specific localization. This is done for all labeled targets within the analysis window and then for all clusters. Again, the clusters used in analysis here were filtered for size as described in the main text. The distance results are then visualized in a standard histogram as shown in Fig.3 B.

For three label analysis, an analysis window of 5× scaling of the area of the original cluster is used. Prior to calculating joint density, the individual co-inhabitance of the marks is measured by going to every localization of either marker and counting the occurrences of the other marker not centered within a 100 nm radius circle (Fig.S5). If there is co-inhabitance of the centered mark and the other labeled target, we then assign a numerical value of 1, if there is not then that localization remains at a base value of zero (Fig. S5). These values are then assigned a distance relative to the original cluster size r0, such that they fit into bins which are expansions upon the original cluster. 50 expansions are used for each cluster, and with normalization of distance to the original cluster size, the co-inhabitance data points are then placed into bins from 0-50 (r/r0). Normalization allows for concatenation of co-inhabitance for a given pairing (RNAPII in H3k27ac and vice versa) since they are all normalized to the cluster size r0. Results for each cluster are combined to give a plot for affinity of one target to the other (see Fig. 4) and then the joint density is given as the combination of each individual plots by taking the counts for each bin and calculating the geometric mean. This is done for each bin from 0 to 5 times the original area. Once this is done, we have the joint density plots as seen in figure 4. These plots are useful due to their ability to highlight enriched areas of both marks to give a better understanding of the distribution of these labeled targets.

### Simulated Distributions Generation

To test the robustness of the algorithms, we generated sample distributions that were spatially distinct. These sample distributions were tested on both image and point-cloud based Euclidean distance measurements as well as on the 3-color point-cloud joint density algorithm. To generate these samples, we sampled according to the distribution requirements. Sampling was done such that the average distribution around all clusters would follow the distribution of interest. The number of samples to be deposited was calculated by finding the average density of the given mark (RNAP II/H3K27ac) in a circular area around the cluster defined by 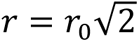 such that the area of the annulus outside the cluster is equal to that inside the cluster with assumed radius *r*_0_. The number of simulated marks deposited would then be given by 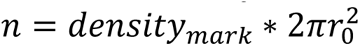. This sampling is relevant for all distributions other than unform where the samples are deposited at uniform positions of 10 nm around the cluster. Samples were deposited around each analysis area around each cluster and representative images of the scatter plots of the resultant distributions for all clusters are shown in Fig. S6A. For image-based algorithm, the samples were binned to the nearest 26 nm pixel and deposited around each identified cluster. Resultant overlays of all samples on one image are shown in Fig. 6B.

### DBSCAN parameter grid search algorithm

Appropriate clustering parameters, epsilon and number of points per cluster, were evaluated by Domain Shape score which considers the size and shape of clusters in an equal manner. We developed this score and performed a parameter grid search to find the optimal epsilon and number of points to accurately capture our targeted structures.

The premise of our scoring involves fitting of an ellipse to the DBSCAN identified clusters and then seeing which parameters optimize our fitting. The optimization is based on a scoring function shown below as well as lower and upper bounds of size which follow biologically relevant sizes identified previously in a publication via Chromatin Scanning Transmission Electron Microscopy^26^.

The general scoring function follows:

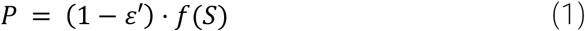

Where Epsilon is the eccentricity of an ellipse, and f(S) is a size function that considers the size of an ellipse (*S* = *πab*) and has the form of:

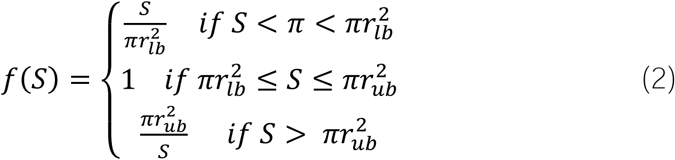

Equation 2 is used to calculate score with equation 1 for all domains. Once that is done, we do a weighted average with weight proportional to size S. The weighted average will further punish the scores and follows the following form:

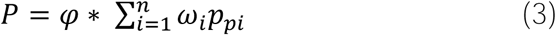

Where *φ* is the usage efficiency that considers the total number of points being used to classify clusters out of all localization points in the dataset. This is used to penalize parameter sets that lead to very large clusters or those that discard many of the clusters as noise. This weighted average will be done for every domain from i=1 to n and will weigh the scores calculated in equation one by their respective weight.

Ellipses were fit where at least three or more peripheral cluster points were considered for circle fitting and five or more points were considered. Normalization of area was done via dividing estimated cluster area by area of a circle of with radius 80 nm. This approach is based on SMLM clustering, however future directions should consider using a ground truth dataset coming from Electron Microscopy or another technique that can resolve domains directly. In this manner, the domain shape score will aim to establish clustering parameters that recapitulate the known size and locations of clusters that have been established *a priori*. Results of grid search can be found in Fig. S3.

Our basis for these analyses is the heterochromatic cluster, which we establish with the standardized DBSCAN algorithm and inform our cluster threshold for size based on prior published identification of chromatin packing domains via electron microscopy^51^. Selecting appropriate parameters for clustering in SMLM data is crucial, as incorrect choices can lead to misleading results. If the epsilon value or the number of points is too high, large chromatin clusters may form, obscuring sub-diffraction chromatin aggregations; too low, and clusters of interest may fragment into smaller, insignificant groups. This parameter optimization is a critical first step, as varying target densities necessitate tailored settings to avoid biasing downstream analyses, especially those relying on cluster peripheries for distance and joint inhabitance studies. While DBSCAN is a useful density-based clustering algorithm, it requires fine-tuning to accurately identify clusters that reflect the biological structure’s size.

### Reagent List

Reagent List:
| Reagent | Components |
| --- | --- |
| Primary Antibodies | H3K9me3 (ab176916, Abcam)<br>H3K27ac (MA5-23516, Thermo Fisher)<br>H3K4me3 (ab213224, Abcam)<br>RNAPII (ab252855, Abcam) |
| Secondary Antibodies | Goat anti Rabbit AF647(A21245, Thermo Fisher)<br>Goat anti Mouse AF568(A11004, Thermo Fisher)<br>Goat anti Rat AF488 (A150157, Thermo Fisher) |
| Blocking Buffer | 3% Bovine Serum Albumin (BSA)<br>0.5% Triton X-100<br>In 1xPBS<br>Modified:<br>Same as Above + 10% Goat Serum |
| Washing Buffer | 0.2% Bovine Serum Albumin (BSA)<br>0.1% Triton X-100<br>In 1X PBS<br>Modified:<br>Same as Above + 1% Goat Serum |

### Detailed Protocol

The 3-color SMLM sample preparation has 3 sequential staining processes for 3 targets.

1. The cells were plated on No. 1 borosilicate bottom eight-well Lab-Tek Chambered cover glass with seeding density of 12.5k. After 48 h, the cells were fixed in 3% paraformaldehyde and 0.1% glutaraldehyde in PBS for 10 min, and then washed with PBS once, quenched with freshly prepared 0.1% sodium borohydride in PBS for 7 min, and rinsed with PBS three times at room temperature.
2. The fixed samples were permeabilized with a blocking buffer (3% bovine serum albumin (BSA), 0.5% Triton X-100 in PBS) for 20 min and then incubated with rabbit anti-H3K9me3 (Abcam) in blocking buffer for a minimum of 2 hours at room temperature and rinsed with a washing buffer (0.2% BSA, 0.1% Triton X-100 in PBS) three times.
3. The fixed samples were further incubated with the corresponding goat secondary antibody–dye conjugates, anti-rabbit AF647 (Thermo Fisher), for 40 min, washed thoroughly with PBS three times at room temperature and stored at 4°C. Upon this step, the staining for the first target (H3K9me3) is finished. The sample can be imaged for single color on heterochromatic target H3K9me3 or incubated overnight in blocking buffer with serum (90% of the blocking buffer mentioned above + 10% goat serum) for the following 2-color and 3-color staining.
4. After overnight blocking step, the sample would go through a similar process as Step 3. But the primary incubation and secondary incubation time are both 1 hour. The blocking buffer will be modified to blocking buffer with serum (90% of the blocking buffer mentioned above + 10% goat serum) and the washing buffer to washing buffer with serum (99% of the blocking buffer mentioned above + 1% goat serum) The primary antibody is rat anti-RNA Polymerase II (Abcam) and the secondary antibody is goat anti-rat AF488 (Abcam). Upon the finish of the RNA Polymerase II staining, the sample can be imaged for 2-color on heterochromatin target H3K9me3 and functional target Polymerase II or incubated overnight in blocking buffer with serum for 3-color staining
5. After overnight blocking step, the sample would go through a similar process as Step 3. But the primary incubation and secondary incubation time are both 1 hour. The blocking buffer will be modified to blocking buffer with serum (90% of the blocking buffer mentioned above + 10% goat serum) and the washing buffer to washing buffer with serum (99% of the blocking buffer mentioned above + 1% goat serum) The primary antibody is mouse anti-H3K27ac (Thermofisher) and the secondary antibody is goat anti-rat AF568 (Abcam). Upon the finish of the RNA Polymerase II staining, the sample can be imaged for 2-color on heterochromatin H3K9me3, functional target Polymerase II and euchromatin target H3K27ac.

