## Supplemental Figures for "Three-color single-molecule localization microscopy in chromatin"

Supplementary Figure 1

A

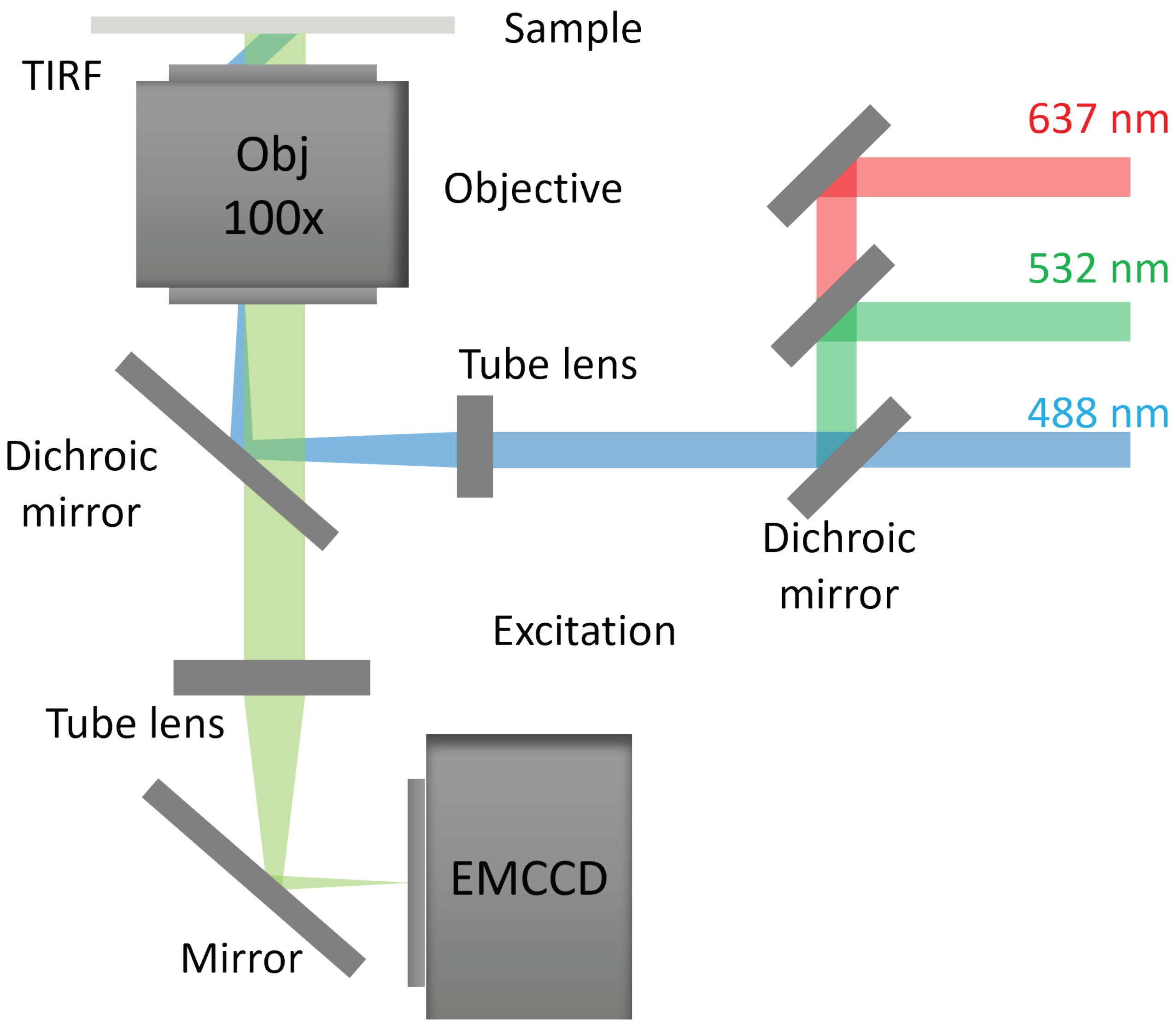

#### Supplementary Figure 2

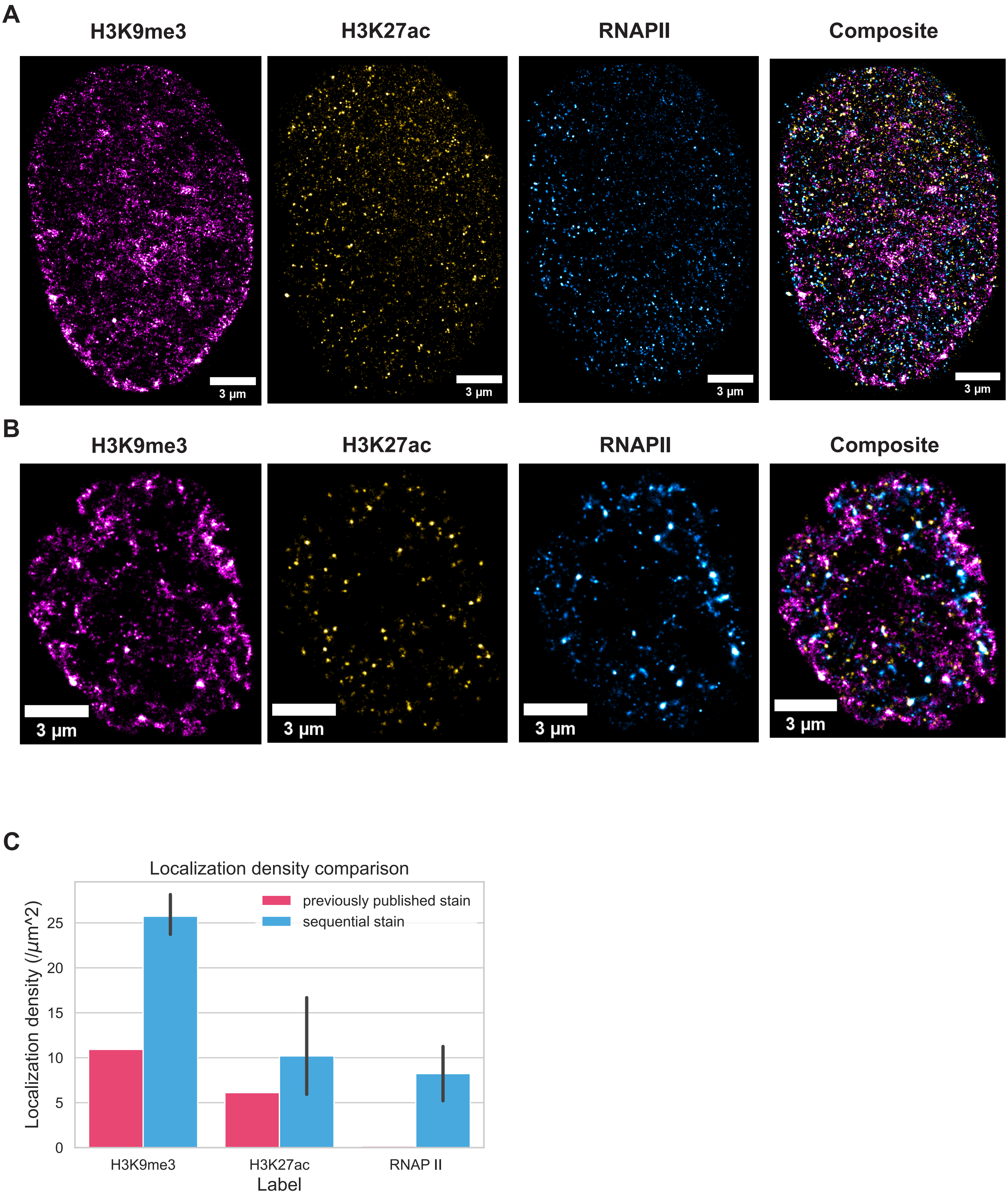

### Supplementary Figure 6

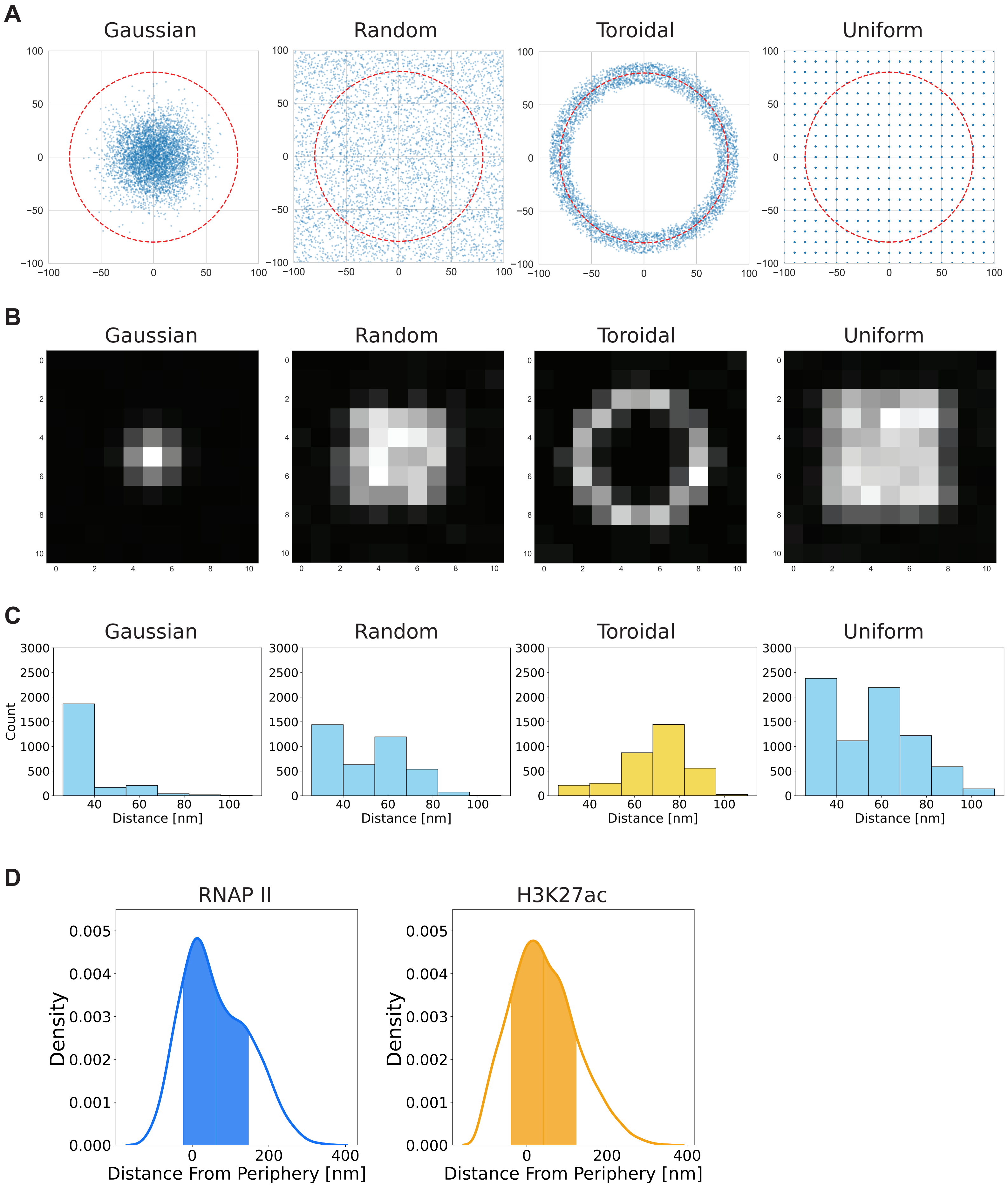

### Supplementary Figure 4

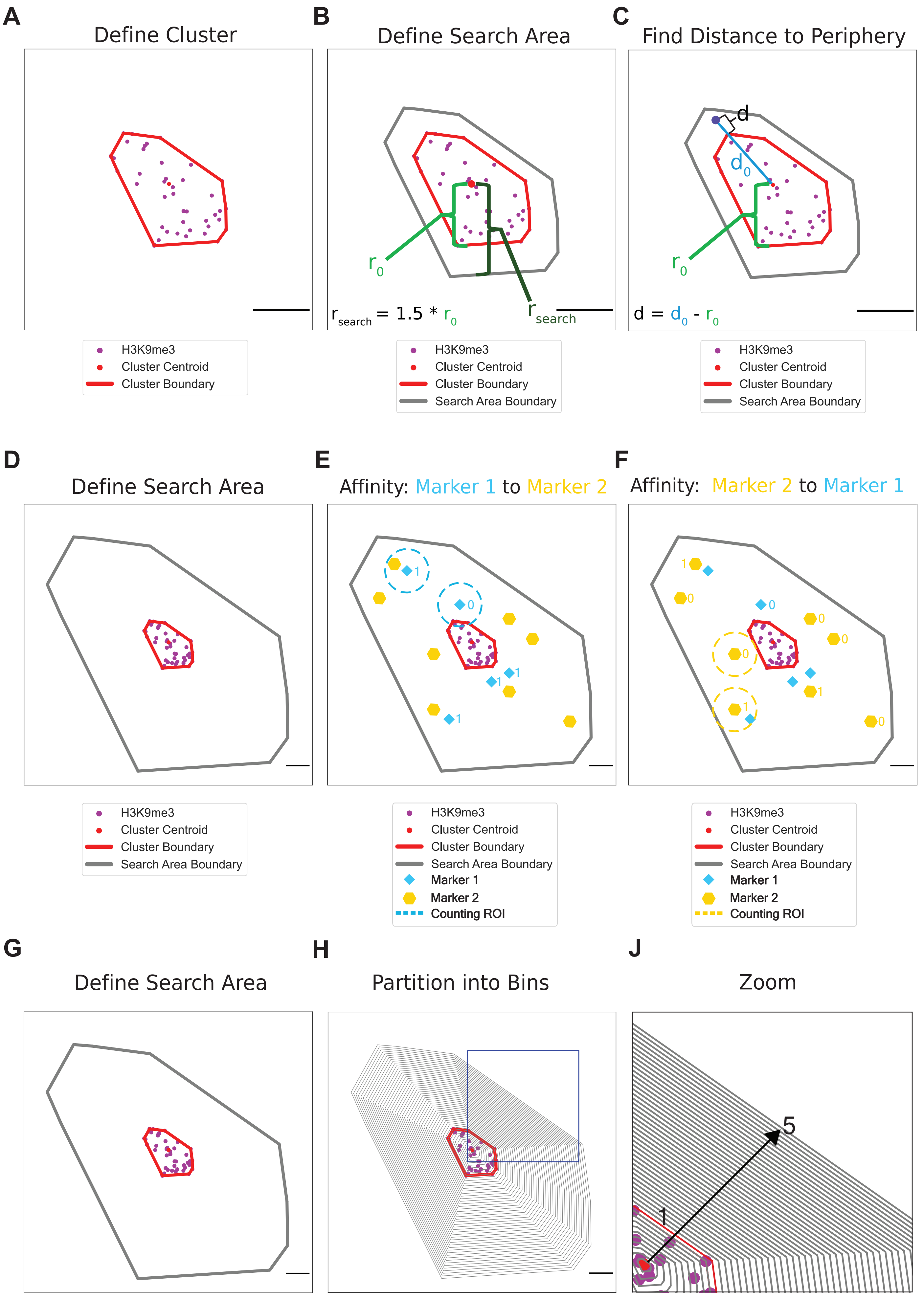

### Supplementary Figure 5

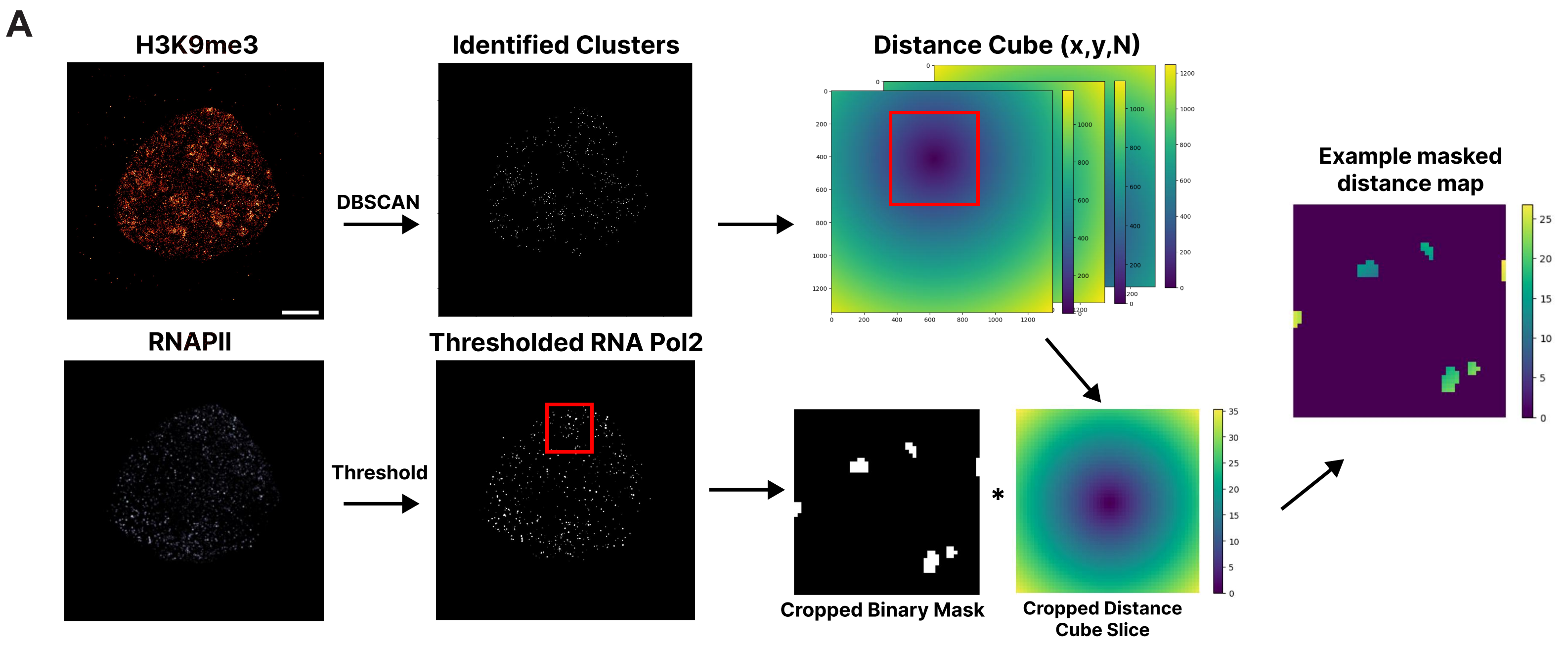


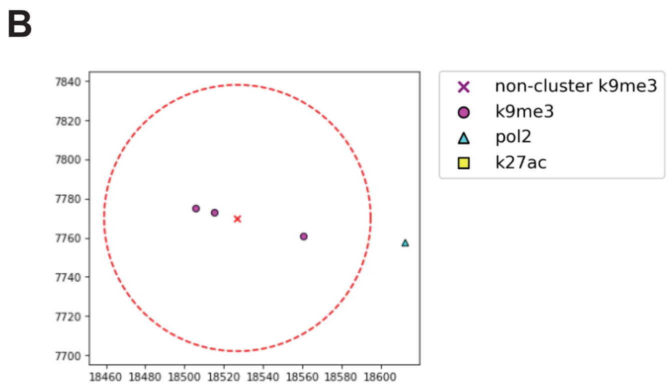

Supplementary Figure 3

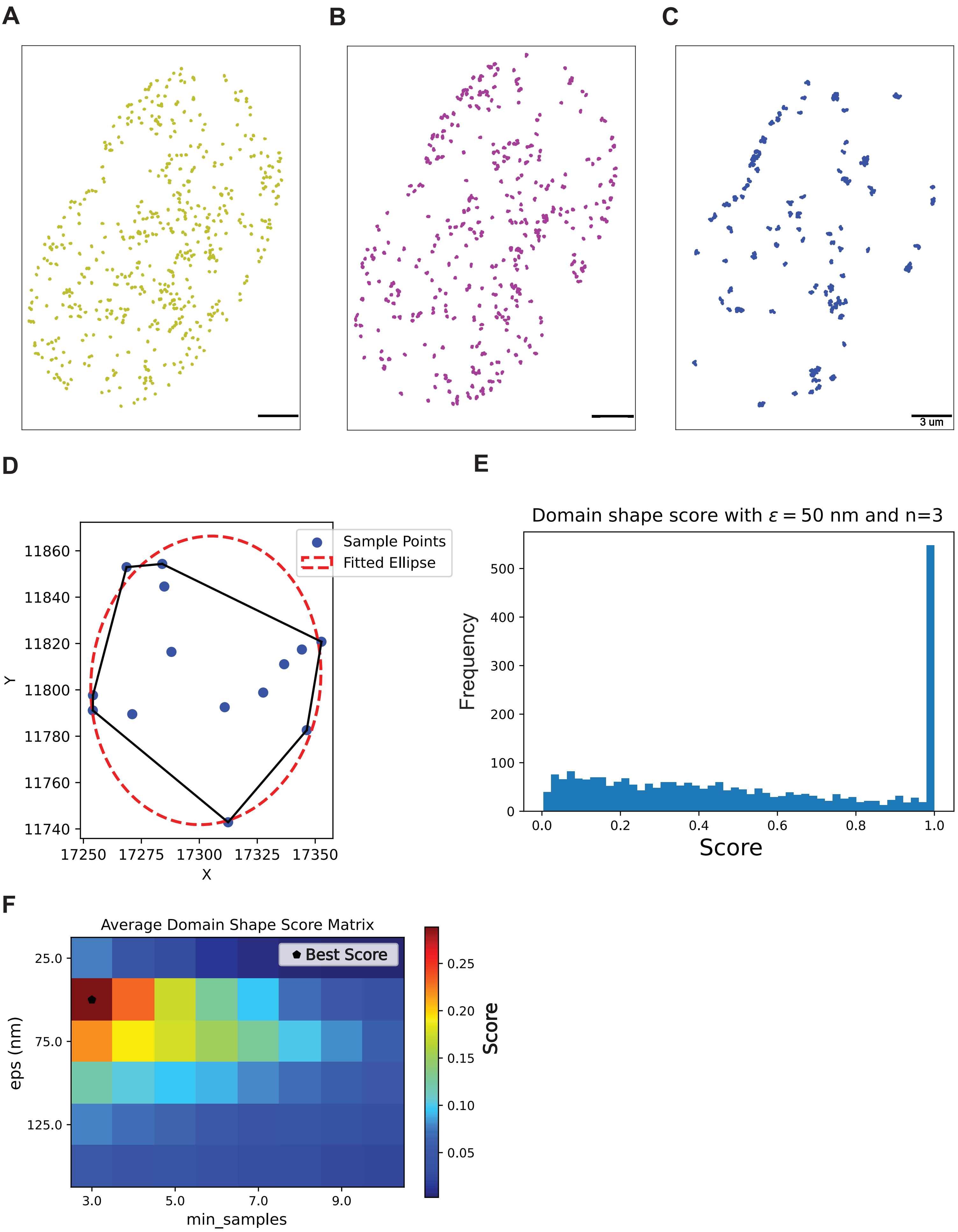
